# ROCK1 regulates insulin secretion from β-cells

**DOI:** 10.1101/2021.06.25.449947

**Authors:** Byung-Jun Sung, Sung-Bin Lim, Jae Hyeon Kim, Won Mo Yang, Rohit N. Kulkarni, Young-Bum Kim, Moon-Kyu Lee

**Author notes:** Corresponding author: Moon Kyu Lee, MD/PhD, Division of Endocrinology and Metabolism, Department of Internal Medicine, Nowon Eulji University Hospital, 68 Hangeulbiseok-ro, Seoul, Korea; Young-Bum Kim, Ph.D., Division of Endocrinology, Diabetes and Metabolism, Beth Israel Deaconess Medical Center, 330 Brookline Avenue, Boston, MA 02215. These authors contributed equally to this work (Young-Bum Kim, Moon-Kyu Lee).

## Abstract

**Objective:** The endocrine pancreatic β-cells play a pivotal role in the maintenance of whole-body glucose homeostasis and its dysregulation is a consistent feature in all forms of diabetes. However, knowledge of intracellular regulators that modulate β-cell function remains incomplete. We investigated the physiological role of ROCK1 in the regulation of insulin secretion and glucose homoeostasis.

**Methods:** Mice lacking ROCK1 in pancreatic β-cells (RIP-Cre; ROCK1^loxP/loxP^, β-ROCK1^-/-^) were studied. Glucose and insulin tolerance tests as well as glucose-stimulated insulin secretion (GSIS) were measured. Insulin secretion response to a direct glucose or pyruvate or pyruvate kinase (PK) activator stimulation in isolated islets from β-ROCK1^-/-^ mice or β-cell lines with knockdown of ROCK1 were also evaluated. Proximity ligation assay was performed to determine the physical interactions between PK and ROCK1.

**Results:** Mice with a deficiency of ROCK1 in pancreatic β-cells exhibited significantly increased blood glucose levels and reduced serum insulin without changes in body weight. Interestingly, β-ROCK1^-/-^ mice displayed progressive impairment of glucose tolerance while maintaining insulin sensitivity mostly due to impaired GSIS. Consistently, GSIS was markedly decreased in ROCK1-deficient islets and ROCK1 knockdown INS-1 cells. Concurrently, ROCK1 blockade led to a significant decrease in intracellular calcium levels, ATP levels, and oxygen consumption rates in isolated islets and INS-1 cells. Treatment of ROCK1-deficient islets or ROCK1 knockdown β-cells either with pyruvate or a PK activator rescued the impaired GSIS. Mechanistically, we observed that ROCK1 binding to PK is greatly enhanced by glucose stimulation in β-cells.

**Conclusions:** Our findings demonstrate that β-cell ROCK1 is essential for glucose-stimulated insulin secretion and maintenance of glucose homeostasis and that ROCK1 acts as an upstream regulator of glycolytic pyruvate kinase signaling.

## 1. INTRODUCTION

Diabetes is a rapidly growing health problem worldwide as evidenced by increasing morbidity and mortality and the economic burden on society [1; 2]. Diabetes is characterized by hyperglycemia, peripheral insulin resistance and impaired β-cell function [3; 4]. Pancreatic β-cells directly contribute to the regulation of systemic glucose balance and by releasing insulin primarily in response to glucose, although other nutrients such as fatty acids and amino acids can also enhance insulin secretion [5; 6]. Impaired glucose-stimulated insulin secretion secondary to defects in signaling pathways has been reported to be a contributing factor for the development of systemic glucose intolerance and the development of overt diabetes [7; 8]. Therefore, identifying molecular mediators that underlie the precise regulation of glucose-induced insulin secretion are of great significance.

Glucose is the principal stimulator of insulin secretion from β-cells acting via the glycolytic pathway [9]. Pyruvate kinase (PK), an enzyme involved in the terminal step of glycolysis, catalyzes the transfer of a phosphate group and converts phosphoenolpyruvate to adenosine diphosphate, to produce pyruvate and ATP [10]. Recent studies demonstrate that activation of PK promotes insulin section during glucose stimulation in INS-1 β-cells and human islets [11]. Furthermore, PK activators can enhance insulin secretion from normal, high-fat diet fed, or Zucker diabetic fatty rats, and diabetic humans [12] indicating the significance of PK in regulating β-cell secretory function.

ROCK1 (Rho-kinase 1; Rho-associated coiled-coil-containing kinase 1) is involved in the pathogenesis of metabolic-related diseases, including hypertension, arteriosclerosis, Alzheimer’s disease, and diabetes [13-15]. Emerging evidence shows that peripheral ROCK1 controls insulin-mediated glucose metabolism and insulin signaling, whereas brain ROCK1 plays a dominant role in regulating feeding behavior and body-weight homeostasis [16-24]. In the liver, ROCK1 is necessary for the development of diet-induced insulin resistance and hepatic steatosis in rodents and humans [25]. Although one study observed that inhibition of ROCK by chemical blockers promotes glucose-stimulated insulin secretion in primary pancreatic β-cells [26], the precise pathways and signaling proteins involved are virtually unexplored. Furthermore, interpretation of this study with inhibitors is limited owing to a lack of ROCK isoform selectivity and incomplete understanding of their respective specificities [27-29].

In the current study, we investigated the role of ROCK1 in regulating glucose metabolism by studying mice lacking ROCK1 in pancreatic β-cells *in vivo* and ROCK1-deficient islets *ex vivo* as well as cultured β-cell lines *in vitro*, with particular emphasis on glucose-stimulated insulin secretion and whole-body glucose homeostasis.

## 2. MATERIALS AND METHODS

### 2.1. Animal care

All animal care and experimental procedures were conducted in accordance with the National Institute of Health’s Guide for the Care and the Use of Laboratory Animals and approved by the Institutional Animal Care and Use Committee of Samsung Medical Center (Seoul, Republic of Korea) and Beth Israel Deaconess Medical Center (Boston, MA). Mice were housed at 22– 24 °C on a 12 h light-dark cycle and allowed *ad libitum* access to standard chow (PicoLab® Rodent Diet 5053, LabDiet, St. Louis, MO) and water.

### 2.2. Generation of RIP-Cre; ROCK1^*loxP/loxP*^ mice

Mice bearing a loxP-flanked ROCK1 allele (ROCK1^loxP/loxP^) were generated and maintained as previously described [22]. Mice lacking ROCK1 in pancreatic β-cells (β-ROCK1^-/-^, RIP-Cre; ROCK1^loxP/loxP^) were generated by breeding ROCK1^loxP/loxP^ mice with RIP-Cre transgenic mice (The Jackson lab, Stock No: 003573). ROCK1^loxP/loxP^ mice were used as controls. All mice were maintained on a mixed genetic background (129Sv and C57BL/6).

### 2.3. Metabolic parameter measurements

Mice were weighed weekly from 5 weeks of age onwards. For daily food intake, 14 week-old males were individually housed for 1 week prior to the start of measurement of food intake. Subsequently, food intake was measured over a 7-day period. Blood was collected via the tail from either randomly-fed or overnight fasted mice. Blood glucose was measured using a glucose meter (Roche, Basel, Switzerland), serum insulin by ELISA (Mercodia, Uppsala, Sweden) and glucagon by ELISA (R&D Systems, Minneapolis, MN, USA).

### 2.4. Oral Glucose tolerance and insulin tolerance test

For the oral glucose tolerance test (OGTT), 8, 20, and 48 week-old males were fasted for 6 h, and blood glucose was measured before and 30, 60, 90, and 120 min after administration of glucose by gavage (2.0 g/kg body weight). For the insulin tolerance test (ITT), 8, 20, and 48 week-old male were fasted for 6 h, and blood glucose was measured before and 15, 30, 60, 90, and 120 min after an intraperitoneal injection of human insulin (0.75 IU/kg body weight: Humulin R, Eli Lilly). Blood glucose was measured using a glucose meter (Roche). The area under the curve for glucose was calculated using the trapezoidal rule for OGTT [30].

### 2.5. Measurement of β-cell numbers and islet insulin content

Pancreas sections (5 µm thick) were immunostained for insulin (guinea pig anti-insulin pig from Abcam, 1:1000, overnight, 4’C, secondary antibody; Alexa Fluor 647-conjugated anti-guinea pig from Invitrogen; 1:500, 1 h room temperature) and nuclei were stained with DAPI (Sigma-Aldrich) in fluorescent mounting medium (Dako). The number of insulin+ β-cells were counted in a random manner by a single observer using a microscope (Olympus Corp.). At least 100 cells were counted per mouse. Insulin levels in islets were measured by ELISA (Mercodia) according to the manufacturer’s instructions.

### 2.6. Pancreatic islet isolation

Pancreases were rapidly dissected after intra-ductal injection from 9-10 week-old male mice and islets were isolated by digesting the pancreas with collagenase P (Roche, Basel, Switzerland) and purified using a Ficoll gradient (Merck Biochrom, Billerica, MA) [31]. After isolation, the islets were cultured for 24 h at 37 °C and size-matched islets were hand-picked using an inverted microscope under sterile conditions.

### 2.7. Perifusion analysis

Isolated islets were preincubated in HEPES-KRB buffer containing 3.3 mM glucose for 30 minutes at 37°C, and then placed in a 0.2 μm syringe filter. The filter was connected to a peristaltic pump and the flow rate was adjusted to 2 or 5 ml/minute. Fractions were serially collected at 5 minute intervals for 30 minutes, 1 minute intervals for 10 minutes, 2 minutes intervals for 50 minutes, and 5 minutes intervals for 30 minutes. Fractions were appropriately diluted and measured for insulin by ELISA (Mercodia).

### 2.8. Glucose-stimulated insulin secretion (GSIS)

The isolated islets or β-cell lines (INS-1 and MIN6 cell) were preincubated with KRB buffer containing 3.3 mM glucose at 37 °C for 120 minutes and then incubated with either 3.3 mM or 16.7 mM glucose in KRB buffer at 37 °C for 60 minutes in the presence or absence of sodium pyruvate (10 mM, Sigma-Aldrich) or TEPP-46 (10 μM, EMD Millipore), or exendin-4 (20 nM, Sigma-Aldrich). Insulin levels in the media were determined by ELISA (Mercodia).

### 2.8. Cell culture for β-cell lines

The INS-1 832/13 cells (gift from C. Newgard PhD, Duke University) were cultured in RPMI-1640 (Gibco) supplemented with 10% FBS and 1% penicillin-streptomycin (Gibco) and HEPES. The MIN6 mouse insulinoma cells were cultured in DMEM (Gibco) supplemented with 15% fetal bovine serum (Thermo Fisher Scientific) and 1% penicillin-streptomycin. The cells were cultured at 37 °C in an atmosphere of 5% CO_2_ and 95% O_2_.

### 2.9. Transfection of INS-1 or MIN6 cells

INS-1 or MIN6 cells were transiently transfected with small interfering RNA (siRNA) using Lipofectamine 2000 (Invitrogen, Carlsbad, CA) according to the manufacturer’s instructions. Cells were used for studies 48 h after transfection. The siRNA sequence for ROCK1 (Bioneer, Daejeon, South Korea) was 5’-UCCAAGUCACAAGCAGACAAGGAUU-3’. Scramble siRNA was used as an experimental control (Bioneer).

### 2.10. Measurement of ATP and calcium levels

The isolated islets or INS-1 cells were starved and incubated with KRB buffer containing 16.7 mM glucose at 37 °C. For measurement of ATP levels, islets or cells were washed with PBS and the bioluminescence reaction was initiated by addition of BacTiter-Glo™ reagent (Promega, Madison, WI) and maintained for 5 min at room temperature. Bioluminescence was determined by a GloMax Multi Microplate Multi Reader (Promega). For measurement of calcium levels, the islets or cells were treated with Fluo-4 direct calcium reagent solution containing 2.5 mM probenecid (Thermo Fisher Scientific) for 60 min at 37°C and washed and then stimulated with 75 mL KCl for 1 min. Bioluminescence was determined by a GloMax Multi Microplate Multi Reader (Promega).

### 2.11. Oxygen consumption rate (OCR) measurements

The isolated islets or INS-1 cells were incubated for 2 h in 2 g/L D-glucose. The OCR was determined using the Seahorse Extracellular Flux (XF-96) analyzer (Seahorse Bioscience). The OCR were calculated by normalizing the protein content for the XF-96 measurement [32].

### 2.12. Pyruvate measurement

The isolated islets or INS-1 cells were preincubated with 3.3 mM glucose for 120 minutes and then incubated with 16.7 mM glucose for 60 minutes. Pyruvate levels in islets and INS-1 cells were measured by Pyruvate Assay Kit (abcam, Cambridge, United Kingdom) according to the manufacturer’s instructions.

### 2.13. Proximity Ligation Assay (PLA)

INS-1 cells were treated with or without fasudil (10 μM) for 1 h and then stimulated with basal (3.3 mM) or high (16.7 mM) glucose for 15 min. Cells were incubated with primary antibodies against ROCK1 (1:100, Cat #: sc-17794, Santa Cruz Biotechnology, Dallas, TX, USA) and pyruvate kinase (1:100, Cat #: ab38237, Abcam) overnight at 4°C. PLA was performed as described previously [33]. PLA was performed using the Duolink® In Situ Detection Reagents Red with Duolink® In Situ PLA probe anti-rabbit PLUS and anti-mouse MINUS (Sigma-Aldrich). The nuclei of cells were stained using Duolink® In Situ Mounting Medium with DAPI (Sigma-Aldrich). Images were captured by a fluorescence microscope (Leica DMi8, Leica) and analyzed by ImageJ (NIH).

### 2.14. Immunoblotting analysis

Tissue lysates (30 μg protein) were resolved by SDS–PAGE and transferred to nitrocellulose membranes (Bio-Rad Laboratories, Hercules, CA). The membranes were incubated with antibodies against ROCK1 (Cat#: sc-17794, Santa Cruz Biotechnology) or ROCK2 (Cat#: sc-5561, Santa Cruz Biotechnology) or β-actin (Cat#: sc-130065, Santa Cruz Biotechnology). The membranes were washed and incubated with secondary antibodies (Cat#: SC-2004 or SC-2005, Santa Cruz Biotechnology). The bands were visualized with SuperSignal Chemiluminescent Substrates (Thermo Fisher Scientific).

### 2.15. Electron microscopy

Pancreas from ROCK1^loxP/loxP^ and β-ROCK1^-/-^ mice fed high glucose were fixed in 2% glutaraldehyde for 12 h. Electron microscopic analysis of docked granules was performed on the pancreas sections and quantitated as described previously [34].

### 2.16. Apoptosis and cell cycle analyses

INS-1 cells were collected for flow cytometry analysis using FITC Annexin V apoptosis detection reagent (BD Biosciences, San Jose, CA). The apoptotic cells were measured by FACS Count™ Flow Cytometer (BD Biosciences). INS-1 cells were stained with propidium iodide (PI, Sigma-Aldrich). Cell cycle analysis was performed using FACS Count™ Flow Cytometer (BD Bioscience).

### 2.17. Statistical Analyses

Data are presented as means ± SEM and individual data points are plotted. Unpaired Student’s ‘t’ tests were used to compare two groups. For comparisons involving more than two groups, one-way analysis of variance (ANOVA) was performed with post-hoc tests and Fisher’s PLSD tests. Repeated measures two-way ANOVA was performed for GTT, ITT, and GSIS. When intervention or interaction (intervention-by-time) was significant by repeated measures two-way ANOVA, post hoc analyses were performed using the SPSS program (SPSS version 18.0, SPSS, Inc., Chicago, IL) for multiple comparisons. All reported p values were two-sided unless otherwise described. Differences were considered significant at *P* < 0.05.

## 3. RESULTS

### 3.1 Selective deletion of ROCK1 in pancreatic β-cells leads to hyperglycemia and impaired glucose-stimulated insulin secretion

We confirmed that ROCK1 expression in pancreatic islets was intact in ROCK1^loxP/loxP^ (control) mice but virtually absent in β-ROCK1^-/-^ mice while its expression in hypothalamus and peripheral tissues was not different between genotypes (Supplementary Figure 1). Furthermore, lack of significant expression of ROCK2 in the islets suggested a lack of compensation between isoforms (Supplementary Figure 1). Body weight (Figure 1A) and daily food intake and accumulated food intake (7 days) were similar between groups (Figure 1B– C). However, β-ROCK1^-/-^ mice displayed hyperglycemia compared with control mice starting at 5 weeks of age, peaked at ∼12 weeks and remained high throughout the 42 week period of the study (Figure 1D). Serum insulin levels in β-ROCK1^-/-^ mice were significantly decreased (∼54%) compared with ROCK1^loxP/loxP^ mice while serum glucagon levels were comparable between groups (Figure 1E–F). Importantly, islet perifusion analysis showed a marked decrease in both the 1^st^ and 2^nd^ phases of insulin secretion in response to glucose stimulation in β-ROCK1^-/-^ mice (Figure 1G). These observations were further confirmed by *ex vivo* statically incubated islet experiments (Figure 1H). However, ROCK1 deletion in pancreatic β-cells had no significant effects on insulin content or β-cell numbers (Figure 1I–J).

**Figure. 1:**
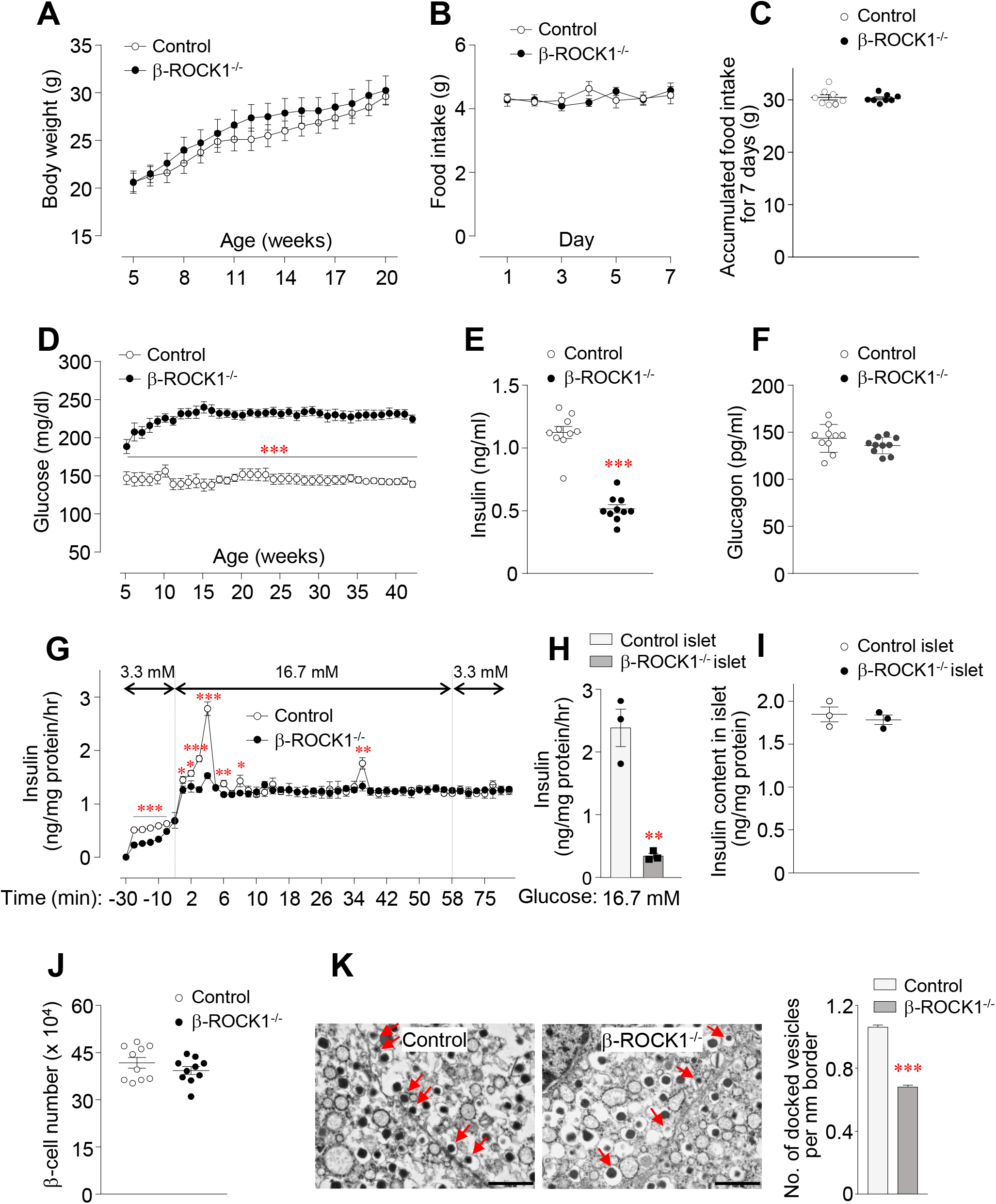
Loss of ROCK1 in pancreatic β-cells leads to hyperglycemia and impaired glucose-stimulated insulin secretion. (A) Body weight (n = 7 for control, n = 7 for β-ROCK1^-/-^), (B) daily food intake (n = 7 for control, n = 7 for β-ROCK1^-/-^), (C) accumulated food intake for 7 days (n = 7 for control, n = 7 for β-ROCK1^-/-^), (D) blood glucose (n = 7 for control, n = 7 for β-ROCK1^-/-^), (E) serum insulin (n = 7 for control, n = 7 for β-ROCK1^-/-^), and (F) serum glucagon (n = 7 for control, n = 7 for β-ROCK1^-/-^) were measured in male control and β-ROCK1^-/-^mice. Serum parameters and fat depots were measured from overnight fasted mice at 9–10 weeks of age. Food intake was measured at 14 weeks of age for 1 week. (G) Perifusion analysis for glucose-stimulated insulin secretion (GSIS) was performed in isolated islets from male control and β-ROCK1^-/-^ mice. (H) GSIS, (I) insulin content, and (J) β-cell number were measured in isolated islets from male control and β-ROCK1^-/-^ mice. (K) Insulin vesicles associated with plasma membrane was assessed in isolated islets from male control and β-ROCK1^-/-^ mice. Arrows indicate membrane-docked vesicles. The scale bar represents 1 μm. Bar graph shows the quantitation of the number of vesicles docked to plasma membrane. All graphs represent means or individual values ± SEM. ^*^*P* <0.05, ^**^*P* <0.01, ^***^*P* <0.001 vs control by two-sided Student’s t-test.

The proximity of insulin granules to the cell surface of β-cells is thought to especially impact the magnitude of the 1^st^ phase insulin release [35; 36]. Electron microscopy analysis showed that the number of secreted granules close to the cell membrane in β-cells of β-ROCK1^-/-^ mice was reduced by 32% compared with β-cells in control mice (Figure 1K, left and right panels), indicating that ROCK1 activation is important in the docking process of insulin granules to the β-cell membrane. Collectively, these results suggest that ROCK1 activation in pancreatic β-cells is involved in the regulation of insulin secretion in vivo.

### 3.2. Mice lacking ROCK1 in pancreatic β-cells are glucose intolerant but not insulin resistant

β-ROCK1^-/-^ mice showed impaired glucose tolerance by age 8 weeks, as revealed by increased area under the glucose curve during OGTT (Figure 2A). While the β-ROCK1^-/-^ mice continued to exhibit glucose intolerance as they aged the intolerance did not worsen (Figure 2B–C). This effect is most likely due to impaired glucose-stimulated insulin secretion and is independent of changes in body weight or insulin sensitivity (Figure 2D–F). Together, these data highlight the necessity of ROCK1 in regulating β-cell function without impacting insulin sensitivity.

**Figure 2:**
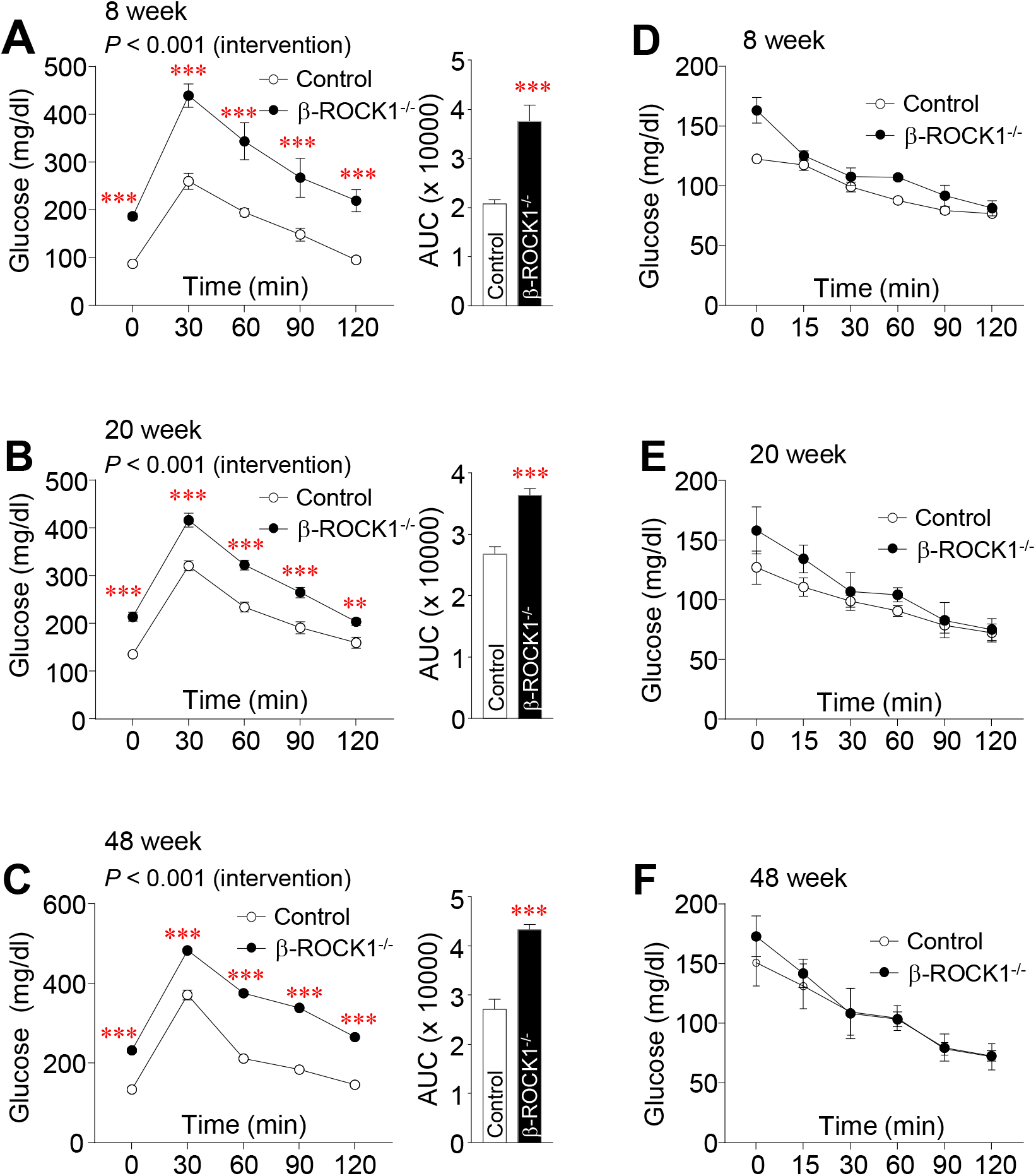
ROCK1 deletion from pancreatic β-cells causes glucose intolerance but not insulin resistance. (A) Oral glucose tolerance test (OGTT) at 8 weeks of age (n = 7 for control, n = 7 for β-ROCK1^-/-^) and (B) OGTT at 20 weeks of age (n = 7 for control, n = 7 for β-ROCK1^-/-^) (C) OGTT at 20 weeks of age (n = 7 for control, n = 7 for β-ROCK1^-/-^) were performed in male control and β- ROCK1^-/-^mice. (D) Insulin tolerance test (ITT) at 8 weeks of age (n = 7 for control, n = 7 for β- ROCK1^-/-^) and (E) ITT at 20 weeks of age (n = 7 for control, n = 7 for β-ROCK1^-/-^) (F) ITT at 20 weeks of age (n = 7 for control, n = 7 for β-ROCK1^-/-^) were performed in male control and β-ROCK1^-/-^mice. Area under the curve (AUC) for GTT was calculated. All graphs represent means ± SEM. *P* values (intervention) for GTT were evaluated by repeated measures two-way ANOVA and *P* values for AUCs were evaluated by two-sided Student’s t-test. ^**^*P* <0.01, ^***^*P* <0.001 vs control mice by repeated measures two-way ANOVA.

### 3.3. Inhibition of ROCK1 impairs glucose-stimulated insulin secretion in INS-1 cells and MIN6 cells

To further determine whether ROCK1 directly regulates insulin secretion in cultured β-cells, we measured GSIS in INS-1 and MIN6 cell lines transfected with ROCK1 siRNA. We confirmed that ROCK1 mRNA levels were significantly reduced by ROCK1 siRNA (Supplementary Figure 2A). Consistent with the *in vivo* results, inhibition of ROCK1 significantly reduced both phases of insulin secretion measured during glucose perifusion in both rodent cell lines (Figure 3A, Supplementary Figure 2B). GSIS was markedly decreased when ROCK1 expression was suppressed (Figure 3B, Supplementary Figure 2C) while insulin content was relatively normal in the cell lines (Figure 3C, Supplementary Figure 2D). These data, combined with the results of in vivo insulin secretion, demonstrate that ROCK1 activation is necessary to regulate glucose-stimulated insulin secretion.

**Figure 3:**
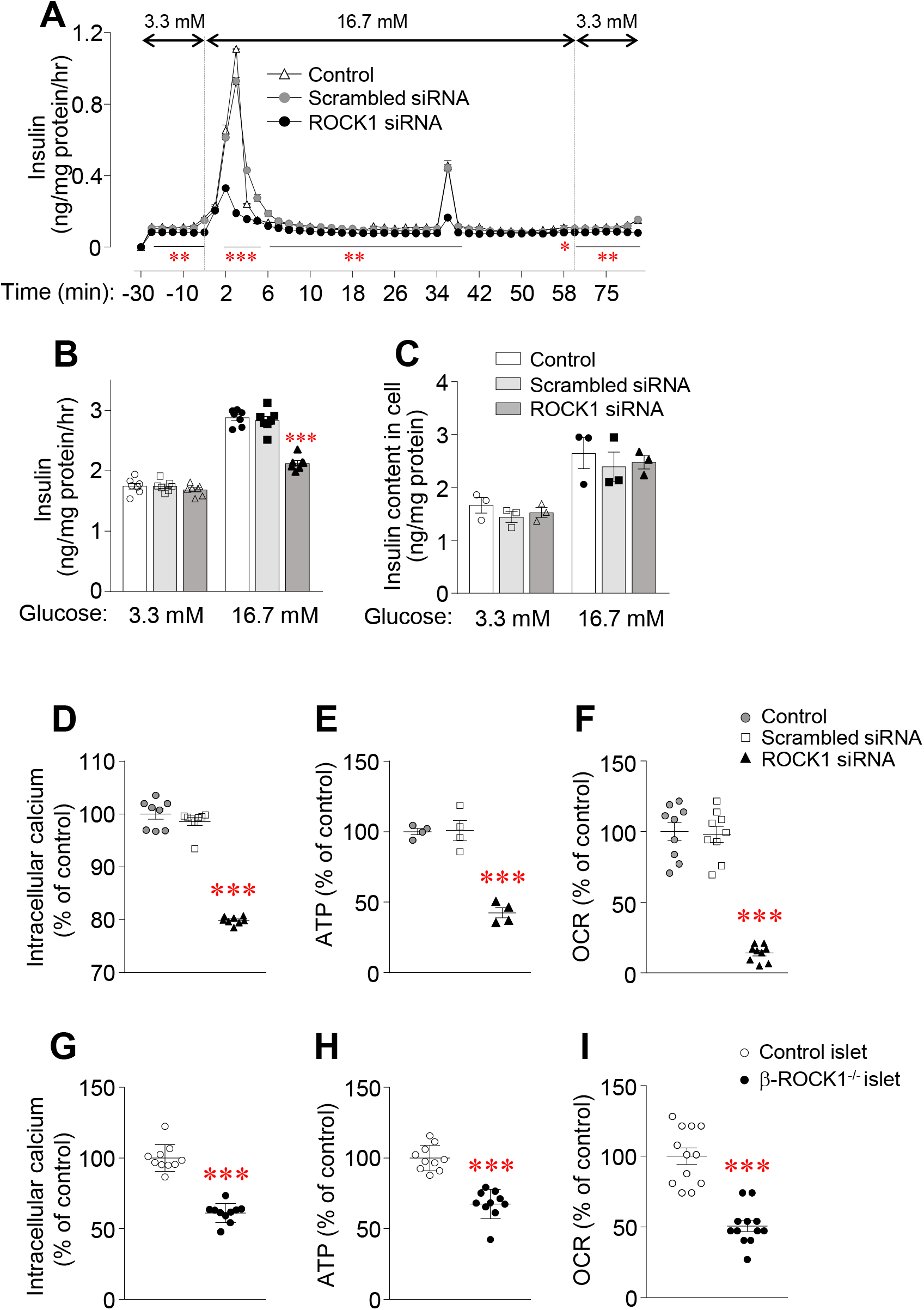
ROCK1 inhibition impairs glucose-stimulated insulin secretion and decreases calcium level, ATP level, and oxygen consumption rate in INS-1 β-cells. (A) Perifusion analysis for glucose-stimulated insulin secretion (GSIS) was performed in ROCK1 knockdown INS-1 cells. (B) GSIS, (C) insulin content, (D) intracellular calcium level, (E) ATP level, and (F) oxygen consumption rate (OCR) were measured in ROCK1 knockdown INS-1 cells. (G) intracellular calcium level, (H) ATP level, and (I) oxygen consumption rate (OCR) were measured in isolated islets from male control and β-ROCK1^-/-^ mice. All graphs represent means or individual values ± SEM. ^*^*P* <0.05, ^**^*P* <0.01, ^***^*P* <0.001 vs scrambled siRNA or control islet by two-sided Student’s t-test.

### 3.4. ROCK1 is involved in the process of insulin secretion

To further investigate the underlying mechanism(s) by which ROCK1 regulates insulin secretion, we measured intracellular Ca^++^ levels, ATP levels and OCR, that are critical events associated with insulin secretion in INS-1 cells. siRNA-mediated inhibition of ROCK1 resulted in a significant decrease in glucose-stimulated intracellular Ca^++^ levels, ATP levels as well as OCR (Figure 3 D–F). Similar findings were observed in islets freshly isolated from β-ROCK1^-/-^ mice (Figure 3 G–I). These data demonstrate that ROCK1 impacts the regulatory machinery involved in insulin secretion in β-cells.

### 3.5 Pyruvate *and* pyruvate kinase (PK) stimulates insulin secretion in the absence of ROCK1

We further determined whether pyruvate, a key intermediate of the glycolysis pathway, is involved in insulin secretion triggered by ROCK1. Pyruvate treatment enhanced the insulin secretion in response to glucose stimulation of INS-1 β-cells transfected with scrambled siRNA. While the insulin secretion was blunted in response to glucose stimualtion in cells with ROCK1 siRNA, the insulin secretion in response to pyruvate was maintained (Figure 4A). Similar observations were found in freshly isolated islets from β-ROCK1^-/-^ mice (Figure 4B). Pyruvate levels were greatly decreased in both INS-1 cell transfected siRNA ROCK1 and islets from β- ROCK1^-/-^ mice (Figure 4C and 4E). Exogenous supplementaion with pyruvate increased insulin secretion 2.9-fold in INS-1 β-cells transfected with siRNA ROCK1 and 3.3-fold in islets from β-ROCK1^-/-^ mice but only 1.3-fold in INS-1 β-cells transfected with scrambled siRNA and 1.5-fold in islets from ccontrol mice (Figure 4D and 4F).

**Figure 4:**
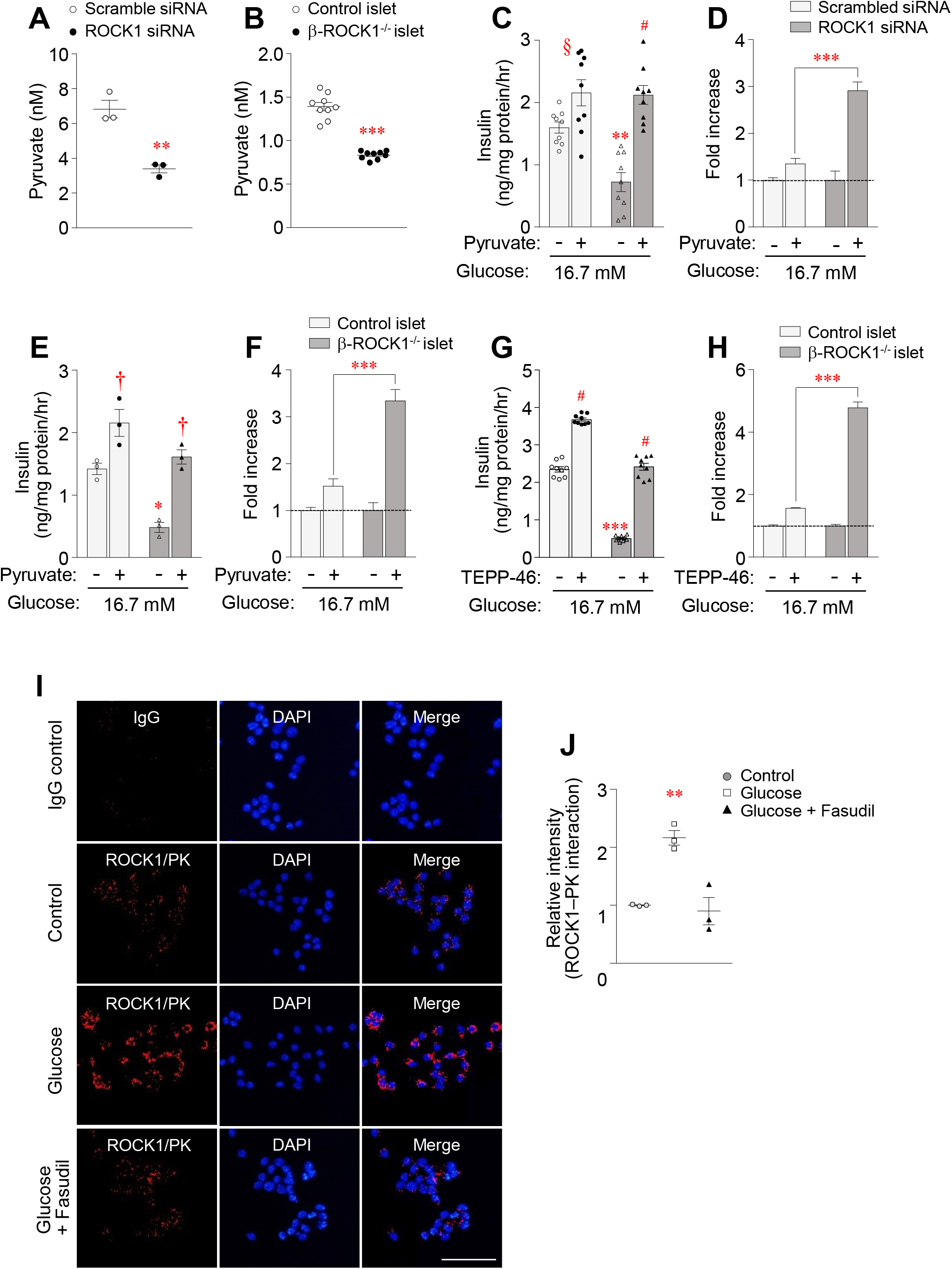
ROCK1 is an upstream regulator of pyruvate kinase (PK) in β-cells. Pyruvate levels were measured in (A) INS-1 cells and (B) islets from control and β-ROCK1^-/-^ mice. Glucose-stimulated insulin secretion (GSIS) was measured in the absence or presence of pyruvate (10 mM) in (C) ROCK1 knockdown INS-1 cells and (E) islets from control and - ROCK1^-/-^ mice. (D, F) Fold increases for GSIS were calculated from C and D, respectively. (E) GSIS was measured in the absence or presence of TEPP-46 (10 μM) in islets from control and -ROCK1^-/-^ mice. (H) Fold increases for GSIS were calculated from G. (I) Proximity ligation assay (PLA) was performed in INS-1 β-cells. Cells were incubated with or without glucose (16.7 mM). Each red spot represents a ROCK1-PK interaction. Nuclei were stained with DAPI (blue). Scale bar represents 25 μm. (J) Image data for ROCK1-PK interaction were quantitated by ImageJ. All graphs represent means or individual values ± SEM. ^*^*P* <0.05, ^**^*P* <0.01, ^***^*P* <0.001 vs scrambled siRNA or control islet, ^§^*P* <0.05, ^†^*P* <0.01, ^#^*P* <0.001 vs no treatment in same group by two-sided Student’s t-test or by repeated measures two-way ANOVA.

Similar to the effects of exogenous pyruvate, the PK activator, TEPP-46, also significantly increased GSIS in freshly isolated islets from control mice (Figure 4G). The effects of the PK activator on enhancing GSIS was evident even in the absence of ROCK1 in isolated islets (Figure 4G). Thus, the PK activator increased insulin secretion 4.8-fold in islets from β-ROCK1^-/-^ mice but only 1.6-fold in in islets from ccontrol mice (Figure 4H). Interestingly, glucagon-like peptide-1 (GLP-1) action was not involved in the regulation of ROCK1-mediated insulin secretion (Supplementary Figure 3A–B). Together, these data suggest that the effects of pyruvate and PK are independent of ROCK1-mediated insulin secretion in β-cells.

### 3.6. ROCK1 interacts with pyruvate kinase

To further test the hypothesis that ROCK1 binds to PK in response to glucose, we undertook proximity ligation assays (PLA), a powerful technology to detect proteins with high specificity and sensitivity [37]. PLA revealed that each red spot represents a ROCK1-PK interaction complex in INS-1 β-cells. As expected, no red spots were detected in IgG control cells. Glucose induced physical interaction between ROCK1 and PK in INS-1 β-cells, as evidenced by a marked increase in the number of red spots. However, this effect was abolished by treatment with the ROCK inhibitor Fasudil (Figure 4I). Quantitative analysis indicated that glucose stimulation increased ROCK1-PK interactions by ∼2 fold over control, and this effect was restored to control levels (Figure 4J). Together, these data suggest that PK physically interacts with ROCK1 in β-cells. The precise dynamic alterations between PK and ROCK1 in disease states warrants further investigation.

## DISCUSSION

The ability of glucose to increase insulin secretion represents a major feature of β-cell function and its dysregulation is a key pathogenic feature in all forms of diabetes [3; 4]. The current study was thus designed to determine the physiological role of ROCK1 in pancreatic β-cells in homeostatic control of insulin secretion in the context of glucose metabolism. Our data clearly suggest that ROCK1 is necessary for the regulation of insulin secretion in pancreatic β-cells during glucose stimulation. This effect is likely mediated through the physical interactions between ROCK1 and PK. Thus, we identify ROCK1 as an important regulator of β-cell metabolism that may lead to new treatment options for diabetes.

A major finding of this study is that deletion of ROCK1 in pancreatic β-cells significantly decreases GSIS in vivo, which ultimately leads to systemic glucose intolerance. These data are confirmed by studies in freshly isolated islets ex vivo and β-cells in vitro. Importantly, these effects are observed when insulin content or β-cell numbers are normal, suggesting that the marked reduction in insulin secretion due to ROCK1 deletion is directly linked to defects in the insulin secretory machinery rather than insulin synthesis and production. This is supported by the reduced number of insulin granules docked at the plasma membrane in β-cells from isolated islets from β-ROCK1^-/-^ mice, which is associated with a decrease in the first phase of insulin release during glucose stimulation. Consistent with this view, previous reports link several proteins such as TRB3 [36], ABCA12 [38] and LKB1 [39] with modulating insulin granules and plasma membrane docking dynamics in the regulation of GSIS. The mechanism(s) underlying insulin granules mobilization could involve in F-actin remodeling in response to glucose stimulation [40]. Given that ROCK regulates actin cytoskeleton reorganization [41; 42], it is conceivable that ROCK1 deletion may inhibit F-actin remodeling to limit the access of insulin granules to the plasma membrane in β-cells.

The glycolytic pathway in pancreatic β-cells involves a cascade of events that break down glucose into pyruvate, producing ATP and nicotinamide adenine dinucleotide (NADH) [43]. Activation of the glycolytic pathway could lead to a significant increase in insulin secretion from β-cells. As expected, exogenous supplementation of β-cells with pyruvate greatly promotes insulin release on a background of glucose stimulation. The fact that pyruvate’s ability to increase insulin secretion during glucose stimulation is enhanced in the absence of ROCK1 suggests that ROCK1-mediated insulin secretory mechanism is linked to the actions of pyruvate. Given that pyruvate levels in islets of β-ROCK1^-/-^ mice were reduced, it is likely that PK, which converts phosphoenolpyruvate and ADP into pyruvate and ATP, is involved in this regulation. The significance of PK activation for the induction of insulin secretion has been recently documented [11; 12]. For example, small molecule activators of PK potently amplify GSIS by switching mitochondria from oxidative phosphorylation to anaplerotic phosphoenolpyruvate biosynthesis [11]. Moreover, PK activation ameliorates GSIS in islets obtained from animals and humans manifesting insulin resistance and type 2 diabetes [12]. In this context, it is notable that we observed that treatment of ROCK1-deficient islets with the PK activator increased glucose-stimulated insulin secretion. Our data point to hypersensitization of β-cells to glucose by pyruvate or PK activator treatment in the absence of ROCK1 that leads to enhanced insulin release. Although the precise underlying mechanism for this phenomenon remains to be elucidated, it is likely that ROCK1-deficient islets are pyruvate-sensitive. Taken together, these novel findings demonstrate that glycolytic PK signaling is linked to ROCK1 action in regulating insulin secretion.

We suggest that ROCK1 is a positive regulator of insulin secretion and that development of small molecules activators for ROCK1 may offer new approaches to treating diabetes.

## AUTHOR CONTRIBUTIONS

The study was designed by B.J.S., Y.B.K., and M.K.L. B.J.S., S.B.L., W.M.Y., and J.H.K. performed in vivo, ex vivo, and in vitro experiments and analyzed data. R.K. provided conceptional advice and contributed to the editing of the manuscript. M.K.L. and Y.B.K. wrote the manuscript with input all other authors. M.K.L. and Y.B.K. are the senior and corresponding author.

## ACKNOWLEDGMENTS

This work was supported by grants from the National Institutes of Health (R01DK111529, R01DK083567, and R01DK123002 to YBK, and R01 067536 to RNK). We thank Aykut Uner for performing statistical analysis.

## Supplementary information

**Supplementary Figure 1:**
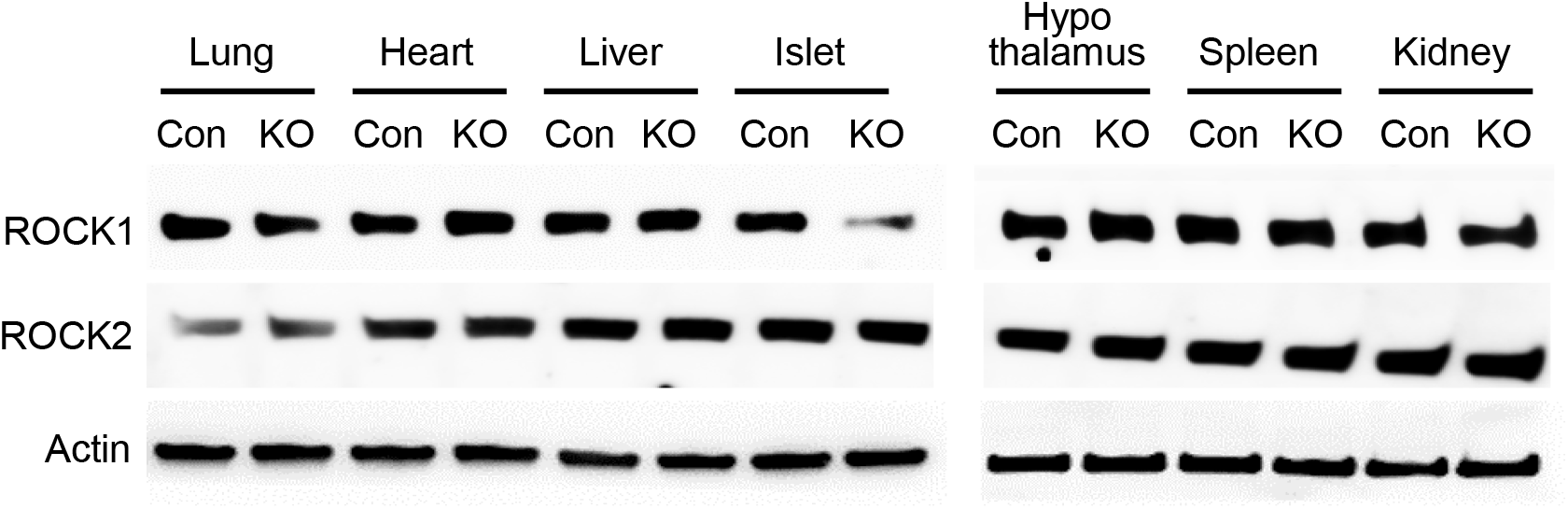
ROCK1 expression in metabolic organs of β-ROCK1^-/-^ mice. Tissue lysates (30 μg) were resolved by SDS-PAGE. ROCK1, ROCK2 or actin was visualized by immunoblotting.

**Supplementary Figure 2:**
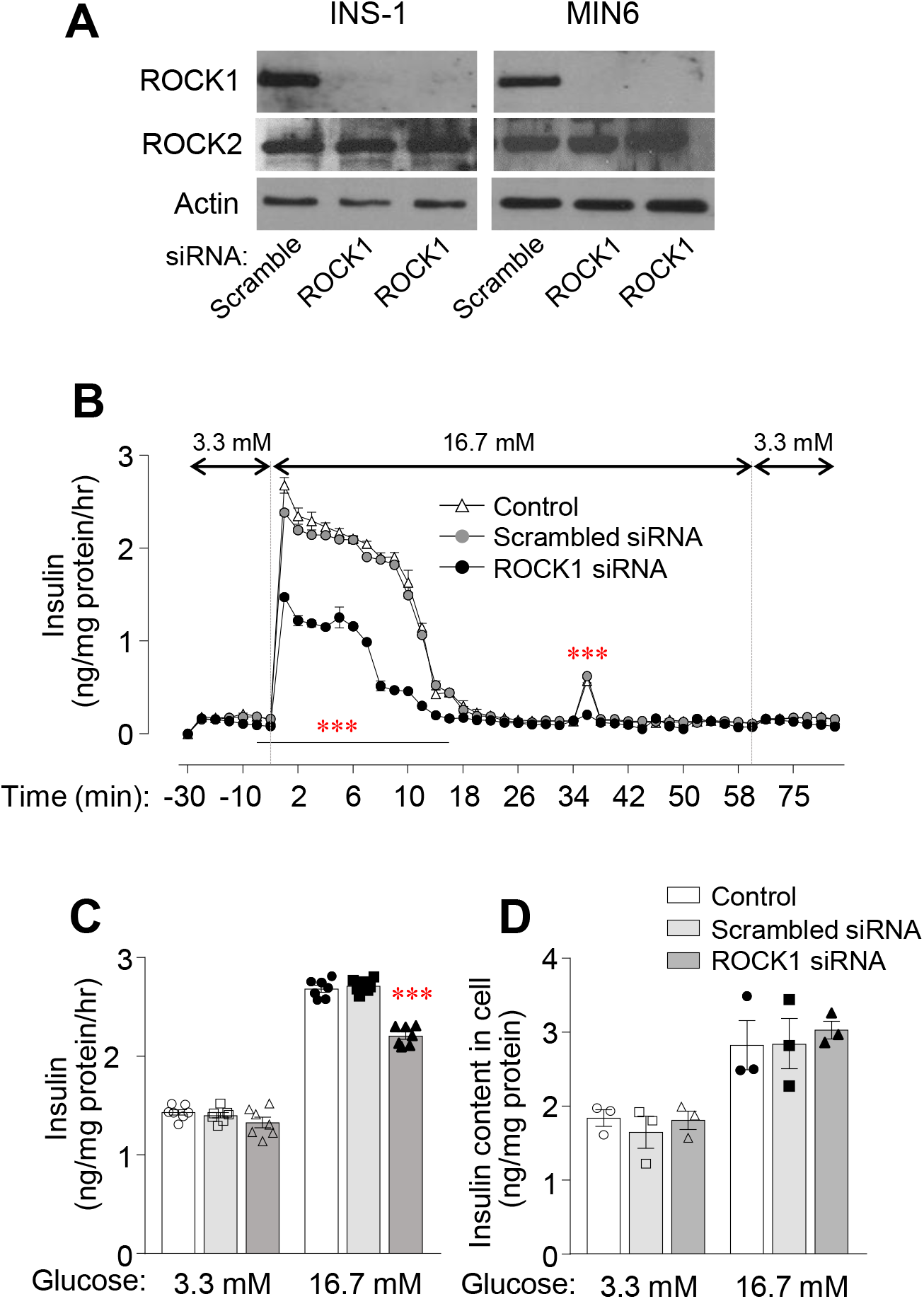
ROCK1 inhibition impairs glucose-stimulated insulin secretion in MIN6 cells. (A) ROCK1 and ROCK2 mRNA levels inROCK1 knockdown INS-1 cells INS-1 and MIN6 cells. mRNA levels of ROCK1 and ROCK2 were measured by reverse transcription PCR (RT-PCR). (B) Perifusion analysis for glucose-stimulated insulin secretion (GSIS) was performed in MIN6 cells. Glucose concentrations during perifusion are indicated. (C) GSIS and (D) insulin content were measured in MIN6 cells. All graphs represent means or individual values ± SEM. ^***^P <0.001 vs scrambled siRNA by ANOVA.

**Supplementary Figure 3:**
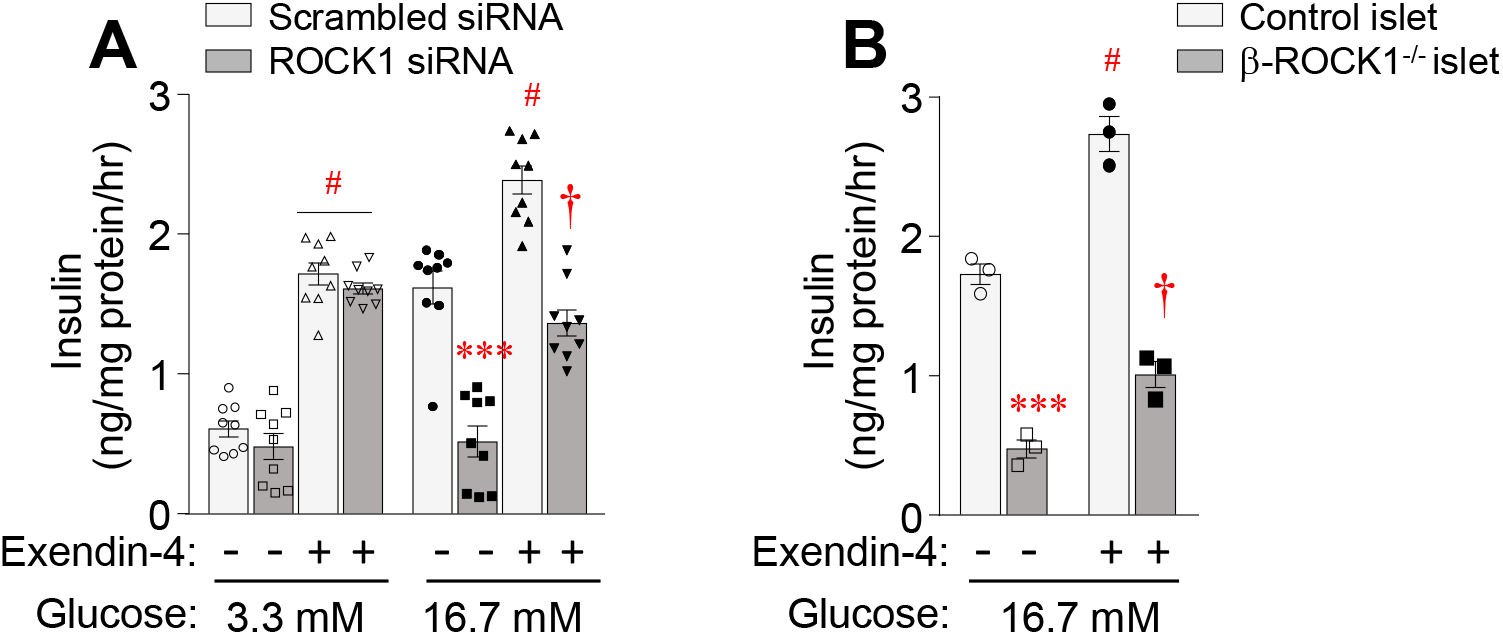
Effect of GLP-1 agonist on glucose-stimulated insulin secretion in INS-1 cells and islets. Glucose-stimulated insulin secretion (GSIS) was measured in the absence or presence of exendin-4 (20 nM) in (A) ROCK1 knockdown INS-1 cells and (B) islets from control and β-ROCK1^-/-^mice. All graphs represent means or individual values ± SEM. ^***^*P* <0.001 vs scrambled siRNA or control islet, ^†^*P* <0.01, ^#^*P* <0.001 vs no treatment in same group by repeated measures two-way ANOVA.

## Reference

[1] Divers, J., Mayer-Davis, E.J., Lawrence, J.M., Isom, S., Dabelea, D., Dolan, L., et al., 2020. Trends in Incidence of Type 1 and Type 2 Diabetes Among Youths - Selected Counties and Indian Reservations, United States, 2002-2015. MMWR Morb Mortal Wkly Rep 69(6):161–165.

[2] Association, A.D., 2018. Economic costs of diabetes in the US in 2017. Diabetes Care 41(5):917–928.

[3] DeFronzo, R.A., 1997. Pathogenesis of type 2 diabetes : metabolic and molecular implications for identifying diabetes. Diabetes Rev 5:177–269.

[4] Taylor, S.I., Accili, D., Imai, Y., 1994. Insulin resistance or insulin deficiency. Which is the primary cause of NIDDM? Diabetes 43(6):735–740.

[5] Bhaswant, M., Poudyal, H., Brown, L., 2015. Mechanisms of enhanced insulin secretion and sensitivity with n-3 unsaturated fatty acids. J Nutr Biochem 26(6):571–584.

[6] de Oliveira, C.A., Latorraca, M.Q., de Mello, M.A., Carneiro, E.M., 2011. Mechanisms of insulin secretion in malnutrition: modulation by amino acids in rodent models. Amino Acids 40(4):1027–1034.

[7] Kulkarni, R.N., Bruning, J.C., Winnay, J.N., Postic, C., Magnuson, M.A., Kahn, C.R., 1999. Tissue-specific knockout of the insulin receptor in pancreatic beta cells creates an insulin secretory defect similar to that in type 2 diabetes. Cell 96(3):329–339.

[8] Withers, D.J., Gutierrez, J.S., Towery, H., Burks, D.J., Ren, J.M., Previs, S., et al., 1998. Disruption of IRS-2 causes type 2 diabetes in mice. Nature 391(6670):900–904.

[9] Prentki, M., Matschinsky, F.M., Madiraju, S.R., 2013. Metabolic signaling in fuel-induced insulin secretion. Cell Metab 18(2):162–185.

[10] Merrins, M.J., Van Dyke, A.R., Mapp, A.K., Rizzo, M.A., Satin, L.S., 2013. Direct measurements of oscillatory glycolysis in pancreatic islet beta-cells using novel fluorescence resonance energy transfer (FRET) biosensors for pyruvate kinase M2 activity. J Biol Chem 288(46):33312–33322.

[11] Lewandowski, S.L., Cardone, R.L., Foster, H.R., Ho, T., Potapenko, E., Poudel, C., et al., 2020. Pyruvate Kinase Controls Signal Strength in the Insulin Secretory Pathway. Cell Metab 32(5):736–750 e735.

[12] Abulizi, A., Stark, R., Cardone, R.L., Lewandowski, S.L., Zhao, Z., Alves, T.C., et al., 2020. Pharmacologic activation of the mitochondrial phosphoenolpyruvate cycle enhances islet function in vivo. bioRxiv doi: https://doi.org/10.1101/2020.02.13.947630.

[13] Hu, E., Lee, D., 2005. Rho kinase as potential therapeutic target for cardiovascular diseases: opportunities and challenges. Expert Opin Ther Targets 9(4):715–736.

[14] Mishra, R.K., Alokam, R., Sriram, D., Yogeeswari, P., 2013. Potential Role of Rho Kinase Inhibitors in Combating Diabetes-Related Complications Including Diabetic Neuropathy-A Review. Curr Diabetes Rev.

[15] Hirooka, Y., Shimokawa, H., 2005. Therapeutic potential of rho-kinase inhibitors in cardiovascular diseases. Am J Cardiovasc Drugs 5(1):31–39.

[16] Begum, N., Sandu, O.A., Ito, M., Lohmann, S.M., Smolenski, A., 2002. Active Rho kinase (ROK-alpha) associates with insulin receptor substrate-1 and inhibits insulin signaling in vascular smooth muscle cells. J Biol Chem 277(8):6214–6222.

[17] Farah, S., Agazie, Y., Ohan, N., Ngsee, J.K., Liu, X.J., 1998. A rho-associated protein kinase, ROKalpha, binds insulin receptor substrate-1 and modulates insulin signaling. J Biol Chem 273(8):4740–4746.

[18] Lee, D.H., Shi, J., Jeoung, N.H., Kim, M.S., Zabolotny, J.M., Lee, S.W., et al., 2009. Targeted disruption of ROCK1 causes insulin resistance in vivo. J Biol Chem 284(18):11776–11780.

[19] Furukawa, N., Ongusaha, P., Jahng, W.J., Araki, K., Choi, C.S., Kim, H.J., et al., 2005. Role of Rho-kinase in regulation of insulin action and glucose homeostasis. Cell Metab 2(2):119–129.

[20] Chun, K.H., Araki, K., Jee, Y., Lee, D.H., Oh, B.C., Huang, H., et al., 2012. Regulation of glucose transport by ROCK1 differs from that of ROCK2 and is controlled by actin polymerization. Endocrinology 153(4):1649–1662.

[21] Chun, K.H., Choi, K.D., Lee, D.H., Jung, Y., Henry, R.R., Ciaraldi, T.P., et al., 2012. In vivo activation of ROCK1 by insulin is impaired in skeletal muscle of humans with type 2 diabetes. Am J Physiol Endocrinol Metab 300(3):E536–542.

[22] Huang, H., Kong, D., Byun, K., Ye, C., Koda, S., Lee, D., et al., 2012. Rho-kinase regulates energy balance by targeting hypothalamic leptin receptor signaling. Nature Neuroscience (15):1391–1398.

[23] Huang, H., Lee, D.H., Zabolotny, J.M., Kim, Y.B., 2013. Metabolic actions of Rho-kinase in periphery and brain. Trends Endocrinol Metab 24(10):506–514.

[24] Huang, H., Lee, S.H., Ye, C., Lima, I.S., Oh, B.C., Lowell, B.B., et al., 2013. ROCK1 in AgRP neurons regulates energy expenditure and locomotor activity in male mice. Endocrinology.

[25] Huang, H., Lee, S.H., Sousa-Lima, I., Kim, S.S., Hwang, W.M., Dagon, Y., et al., 2018. Rho-kinase/AMPK axis regulates hepatic lipogenesis during overnutrition. J Clin Invest 128(12):5335–5350.

[26] Hammar, E., Tomas, A., Bosco, D., Halban, P.A., 2009. Role of the Rho-ROCK (Rho-associated kinase) signaling pathway in the regulation of pancreatic beta-cell function. Endocrinology 150(5):2072–2079.

[27] Ono-Saito, N., Niki, I., Hidaka, H., 1999. H-series protein kinase inhibitors and potential clinical applications. Pharmacol Ther 82(2-3):123–131.

[28] Narumiya, S., Ishizaki, T., Uehata, M., 2000. Use and properties of ROCK-specific inhibitor Y-27632. Methods Enzymol 325:273–284.

[29] Diep, D.T.V., Hong, K., Khun, T., Zheng, M., Ul-Haq, A., Jun, H.S., et al., 2018. Anti-adipogenic effects of KD025 (SLx-2119), a ROCK2-specific inhibitor, in 3T3-L1 cells. Sci Rep 8(1):2477.

[30] Alquier, T., Poitout, V., 2018. Considerations and guidelines for mouse metabolic phenotyping in diabetes research. Diabetologia 61(3):526–538.

[31] Lacy, P.E., Kostianovsky, M., 1967. Method for the isolation of intact islets of Langerhans from the rat pancreas. Diabetes 16(1):35–39.

[32] Lamb, R., Fiorillo, M., Chadwick, A., Ozsvari, B., Reeves, K.J., Smith, D.L., et al., 2015. Doxycycline down-regulates DNA-PK and radiosensitizes tumor initiating cells: Implications for more effective radiation therapy. Oncotarget 6(16):14005–14025.

[33] Seo, J.A., Kang, M.C., Yang, W.M., Hwang, W.M., Kim, S.S., Hong, S.H., et al., 2020. Apolipoprotein J is a hepatokine regulating muscle glucose metabolism and insulin sensitivity. Nat Commun 11(1):2024.

[34] Gomi, H., Mizutani, S., Kasai, K., Itohara, S., Izumi, T., 2005. Granuphilin molecularly docks insulin granules to the fusion machinery. J Cell Biol 171(1):99–109.

[35] Rorsman, P., Renstrom, E., 2003. Insulin granule dynamics in pancreatic beta cells. Diabetologia 46(8):1029–1045.

[36] Liew, C.W., Bochenski, J., Kawamori, D., Hu, J., Leech, C.A., Wanic, K., et al., 2010. The pseudokinase tribbles homolog 3 interacts with ATF4 to negatively regulate insulin exocytosis in human and mouse beta cells. J Clin Invest 120(8):2876–2888.

[37] Soderberg, O., Gullberg, M., Jarvius, M., Ridderstrale, K., Leuchowius, K.J., Jarvius, J., et al., 2006. Direct observation of individual endogenous protein complexes in situ by proximity ligation. Nat Methods 3(12):995–1000.

[38] Ursino, G.M., Fu, Y., Cottle, D.L., Mukhamedova, N., Jones, L.K., Low, H., et al., 2020. ABCA12 regulates insulin secretion from beta-cells. EMBO Rep 21(3):e48692.

[39] Fu, A., Robitaille, K., Faubert, B., Reeks, C., Dai, X.Q., Hardy, A.B., et al., 2015. LKB1 couples glucose metabolism to insulin secretion in mice. Diabetologia 58(7):1513–1522.

[40] Kalwat, M.A., Thurmond, D.C., 2013. Signaling mechanisms of glucose-induced F-actin remodeling in pancreatic islet beta cells. Exp Mol Med 45:e37.

[41] Iizuka, M., Kimura, K., Wang, S., Kato, K., Amano, M., Kaibuchi, K., et al., 2012. Distinct distribution and localization of Rho-kinase in mouse epithelial, muscle and neural tissues. Cell Struct Funct 37(2):155–175.

[42] Kawano, Y., Fukata, Y., Oshiro, N., Amano, M., Nakamura, T., Ito, M., et al., 1999. Phosphorylation of myosin-binding subunit (MBS) of myosin phosphatase by Rho-kinase in vivo. J Cell Biol 147(5):1023–1038.

[43] Sugden, M.C., Holness, M.J., 2011. The pyruvate carboxylase-pyruvate dehydrogenase axis in islet pyruvate metabolism: Going round in circles? Islets 3(6):302–319.

